# The UCSC SARS-CoV-2 Genome Browser

**DOI:** 10.1101/2020.05.04.075945

**Authors:** Jason D. Fernandes, Angie S. Hinrichs, Hiram Clawson, Jairo Navarro Gonzalez, Brian T. Lee, Luis R. Nassar, Brian J. Raney, Kate R. Rosenbloom, Santrupti Nerli, Arjun Rao, Daniel Schmelter, Ann S. Zweig, Todd M. Lowe, Manuel Ares, Russ Corbet-Detig, W. James Kent, David Haussler, Maximilian Haeussler

**Affiliations:** Genomics Institute, University of California Santa Cruz, Santa Cruz, CA, 95064, USA; Howard Hughes Medical Institute, University of California Santa Cruz, Santa Cruz, CA, 95064, USA; ImmunoX Initiative, University of California San Francisco, San Francisco, CA 94143, USA; Molecular, Cell and Developmental Biology, University of California Santa Cruz, Santa Cruz, CA, 95064, USA; Center for Molecular Biology of RNA, University of California Santa Cruz, Santa Cruz, CA, 95064, USA

## Abstract

**Background:** Researchers are generating molecular data pertaining to the SARS-CoV-2 RNA genome and its proteins at an unprecedented rate during the COVID-19 pandemic. As a result, there is a critical need for rapid and continuously updated access to the latest molecular data in a format in which all data can be quickly cross-referenced and compared. We adapted our genome browser visualization tool to the viral genome for this purpose. Molecular data, curated from published studies or from database submissions, are mapped to the viral genome and grouped together into “annotation tracks” where they can be visualized along the linear map of the viral genome sequence and programmatically downloaded in standard format for analysis.

**Results:** The UCSC Genome Browser for SARS-CoV-2 (https://genome.ucsc.edu/covid19.html) provides continuously updated access to the mutations in the many thousands of SARS-CoV-2 genomes deposited in GISAID and the international nucleotide sequencing databases, displayed alongside phylogenetic trees. These data are augmented with alignments of bat, pangolin, and other animal and human coronavirus genomes, including per-base evolutionary rate analysis. All available annotations are cross-referenced on the virus genome, including those from major databases (PDB, RFAM, IEDB, UniProt) as well as up-to-date individual results from preprints. Annotated data include predicted and validated immune epitopes, promising antibodies, RT-PCR and sequencing primers, CRISPR guides (from research, diagnostics, vaccines, and therapies), and points of interaction between human and viral genes. As a community resource, any user can add manual annotations which are quality checked and shared publicly on the browser the next day.

**Conclusions:** We invite all investigators to contribute additional data and annotations to this resource to accelerate research and development activities globally. Contact us at with data suggestions or requests for support for adding data. Rapid sharing of data will accelerate SARS-CoV-2 research, especially when researchers take time to integrate their data with those from other labs on a widely-used community browser platform with standardized machine-readable data formats, such as the SARS-CoV-2 Genome Browser.

## Introduction

The University of California, Santa Cruz (UCSC) Human Genome Browser (Kent et al., 2002) is a web-based, interactive viewer for the human and other vertebrate genome sequences featuring research data, clinical molecular data, annotations, and sequence alignments that has been used for almost 20 years by hundreds of thousands of biomedical researchers and cited in more than 37,500 scientific papers. To address the current COVID-19 epidemic, we have built a similar browser for the SARS-CoV-2 reference genome (NC_045512v2, wuhCor1). The purpose of this report is to introduce this tool to the international community racing to understand the details of the virus, its evolution, its mechanisms of action in human cells, and its immunological and molecular vulnerabilities.

### A brief introduction to the Genome Browser

The SARS-CoV-2 browser displays the reference nucleotide sequence of the viral genome, and provides an intuitive way to visualize annotations or data on specific parts of the genome (Figure 1). The genome sequence is shown from left to right, 5’ to 3’, as an image with the label “NC_045512.2”, which is the NCBI/INSDC accession ID for the reference sequence. This reference sequence is the RNA genome isolated from one of the first cases in Wuhan and is known as “Severe acute respiratory syndrome coronavirus 2 isolate Wuhan-Hu-1, complete genome” (Wu et al., 2020). This sequence is widely used as a standard reference, and, because of its early identification, it is used as the root genome in phylogenetic trees produced by Nextstrain (Hadfield et al., 2018), The COG-UK (Rambaut et al., 2020) and the China National Center for Bioinformation project (Zhao et al., 2020). Above this image are the **Navigation controls.** Buttons in the navigation controls allow the user to move left and right along the genomic sequence and zoom in and out by various factors. Clicking the zoom factor “base” zooms in to show the nucleotide bases in the selected region. The **Position Bar** shows the current region of the genome being viewed via a red box, and the **Search Box** allows a user to manually enter a specific position to view. Coordinates in the Position Bar should be entered with the prefix NC_045512v2 (the NCBI RefSeq accession of this genome) followed by the stop and start nucleotide numbers (e.g. “NC_045512v2:25,341-25,401”). Users can also directly enter a nucleotide sequence, or the name of a gene (corresponding to the NCBI/INSDC annotation), to go directly to that region of the genome.

**Figure 1:**
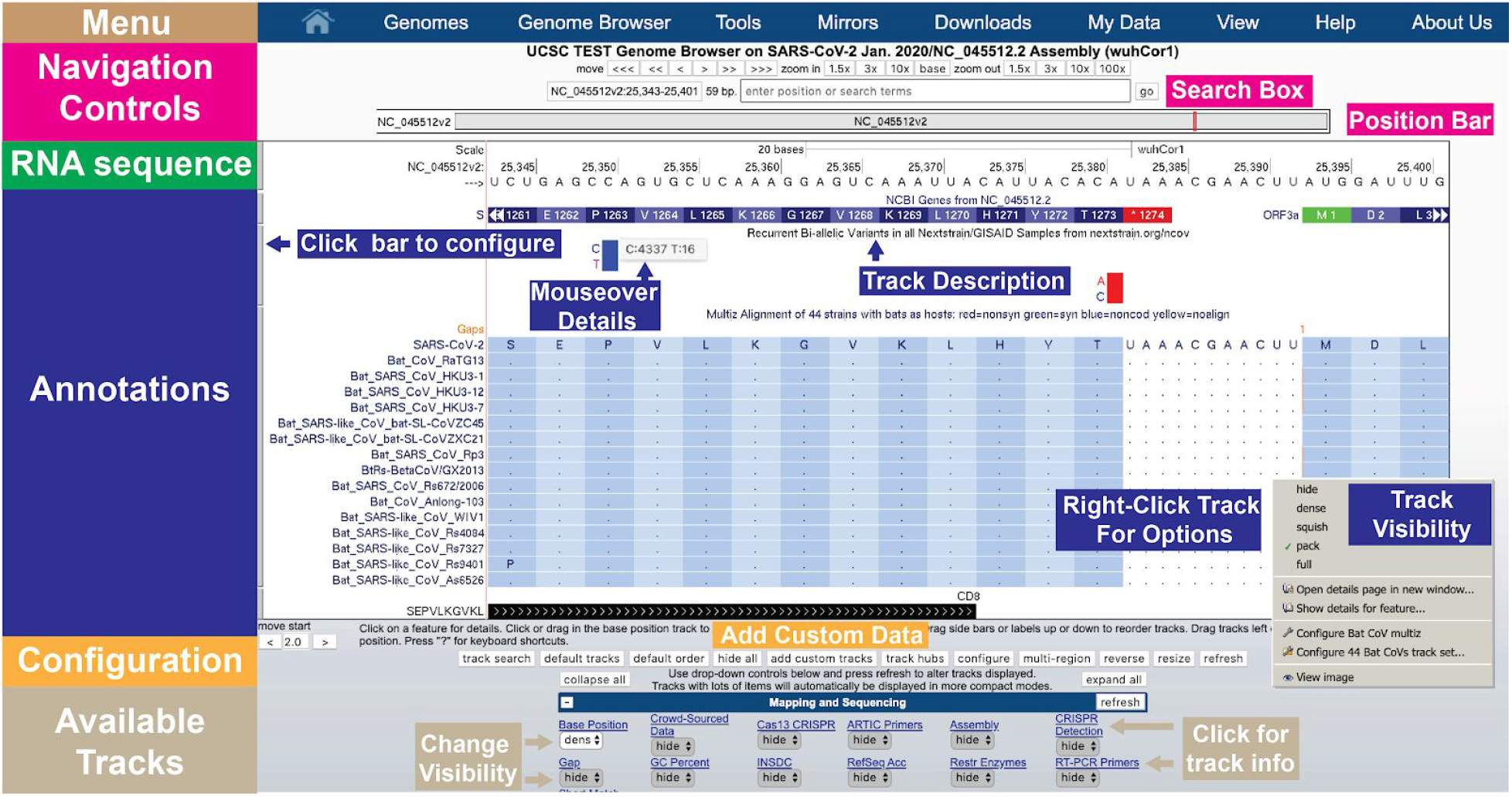
A quick overview of the UCSC Genome Browser user interface structure. Navigation controls at the top allow users to move left and right and to zoom. The Position Bar shows a highlighted red box illustrating the current portion of the genome being viewed. The Search Box allows users to search for particular features or to move to exact genomic coordinates. The RNA sequence is only shown when sufficiently zoomed in. Annotations are shown for data tracks that have been set to visible. Here, the NCBI Genes track shows the annotation of the end of the Spike (S) protein and the start of ORF3a, as well as the amino acid translation of their codons. Below that a track showing recurrent SARS-CoV-2 variants that have been observed around the world as reported by nextstrain.org. Bar graphs show the frequency of each allele and mouseover gives the counts of each allele. The next track (Bat CoV multiz) shows a multiple alignment of 44 bat coronaviruses aligned to the reference. Overall these viruses align well to this region of SARS-CoV-2 (the dot means the amino acid is identical) although one non-synonymous substitution (S1261P) is observed in one virus, Rs9401. The final track shows a CD8 positive epitope from IEDB (see Figure 7 for additional details). Tracks can be configured with a right-click or alternatively by clicking on their name near the bottom of the page. Only 12 of the 48 currently available track configuration buttons are shown in this figure due to space limitations. Custom data tracks generated by users can be added directly via the “add custom tracks” button. Additional options can be set via the Menu bar at the top (e.g. the “View” menu allows additional changes to the browser window). (Live interactive session for this figure: http://genome.ucsc.edu/s/SARS_CoV2/Figure1)

Genome annotations are shown underneath the Position Bar. At the top of the annotations the genome sequence is shown (when zoomed in) as an **RNA Sequence.** At the time of this writing there are 48 tracks with different kinds of molecular information of which 11 are shown by default. The small **grey buttons** on the left of the image are the **track configuration** buttons which allow users to configure the way track information is displayed. Alternatively, users can also **right-click** to configure the track.

Under the genome sequence on the image, **gene and protein annotations** are shown by default as blue rectangles, with the direction of transcription indicated by small arrows. If activated, additional tracks displaying **alignments** and **variants** are shown underneath.

Under the annotations display, various buttons allow users to **reset** the currently shown tracks back to the defaults. There are two buttons, **add custom tracks** and **track hubs** (Raney et al., 2014), that allow experienced users to **add their own data**. The **configure**button allows users to make viewing adjustments (e.g. increase the font size). The **reverse** button reverse complements the display to show the antisense sequence from 3’ to 5’ and the **resize** button fits the image to the current screen size.

Below the buttons, the **track list** shows all available tracks (the first 12 are shown in Figure 1). In order to provide a compact display, most tracks are hidden by default. For example, in Figure 1, only 4 of the more than 50 currently available tracks are visible. Hovering the mouse over the title of a track for a few seconds will reveal a longer description of the track data. Clicking the title of the track will show a full description and configuration page for that track data. On this page, or in the track list, tracks can be set to one of the four visibility modes: **dense**, **squish**, **pack** and **full** (Figure 2). Depending on the track, different viewing modes may emphasize features of the data not immediately apparent in other modes (e.g. in Figure 2 individual ORFs are hidden in “dense”, and names are hidden in “squish”). In general, setting a track to “pack” is a good starting point to explore the data.

**Figure 2:**
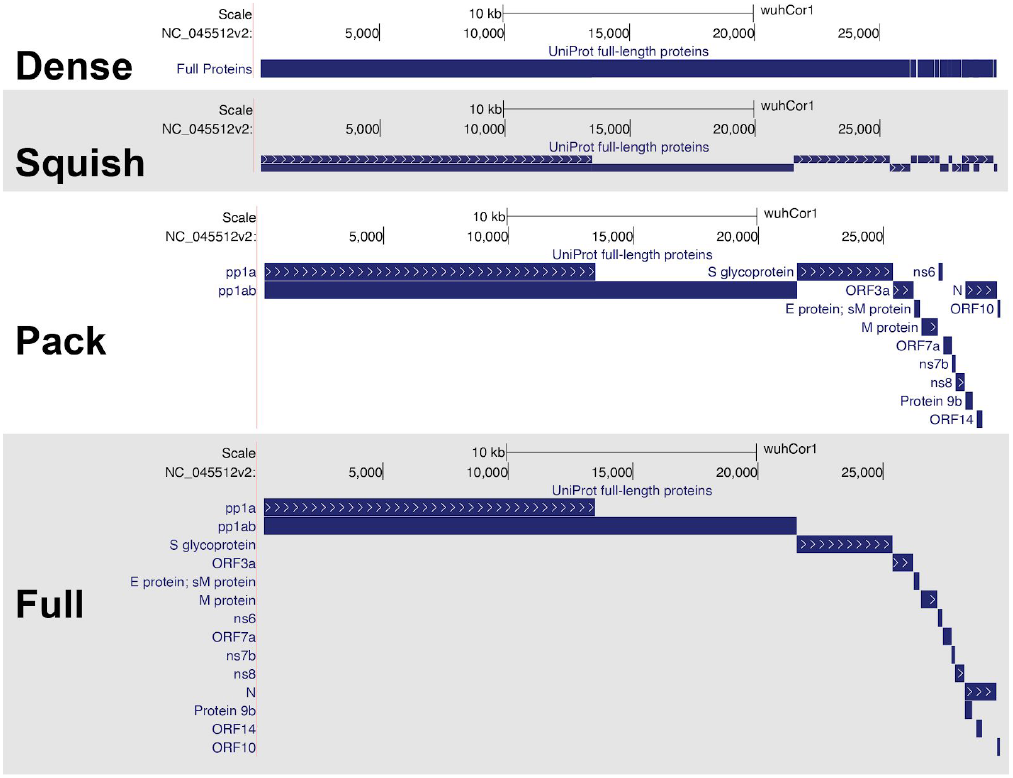
The four visibility modes of annotation tracks. Four different ways to display the protein products of the viral genome. Shown here are the “UniProt full-length proteins” track from the UniProt track collection. “Dense” mode shows a single line highlighting any base that carries an annotation, while “squish”, “pack” and “full” expand the annotations in more detail but use more screen space. (Live Interactive Session: http://genome.ucsc.edu/s/SARS_CoV2/Figure2)

At the top of the page is the **Menu** bar, which contains a few tools helpful for working with the genome sequence:

**Tools > Blat**: Blat (Kent, 2002) is the “BLAST-like alignment tool” that allows users to quickly find the coordinates of a short nucleotide or protein sequence encoded within the genome.
**Tools > Table Browser:** The Table Browser is a simple interface to download the data stored within the annotation tracks. The annotations can be downloaded as spreadsheet tables, or in other common genomics formats (e.g. BED or GFF). Within the Table Browser, users can also perform basic analysis such as intersecting different tracks of data and downloading the resulting files or displaying them as tracks within the genome browser. For API or bulk data access, see “Data Access” below.
**My Data > My Sessions:** Save the current view (“session) to a link that can be shared with others. Then other users can explore this session interactively and make their own track adjustments without affecting the original session shared by the link.
**View > PDF/PS:** Download current view to a PDF or EPS. These files can be used to make high resolution figures for publications and presentations.
**View > DNA:** Download the current genome sequence that is visible within the browser window. Note that although the viral sequence is viewable as RNA (with U instead of T) the downloaded sequence will use the DNA alphabet (T instead of U).

### Genomic Organization of SARS-CoV-2

The reference SARS-CoV-2 genome is a single strand of 29,903 RNA nucleotides, yet it, like many viral genomes, encodes a surprising amount of molecular complexity, generating ~10 canonical RNA transcripts, ~14 ORFs, and ~29 proteins. This complex organization has several features that are atypical for standard genomic analyses and therefore a careful understanding of genomic annotations and viral nomenclature is necessary to accurately interpret the data on the genome browser.

### The SARS-CoV-2 transcriptome

Like other coronaviruses, the SARS-CoV-2 is a single-stranded positive-sense RNA, and shares many features with the mRNA of most human genes, including 5’ and 3’ UTRs as well as a 3’-poly-A tail (Chen et al., 2020). In infected cells, the first gene (orf1a/orf1ab) is directly translated and cleaved into proteins that form the replication-transcription complexes (RTCs) (Nakagawa et al., 2016). These RTCs then use the same positive-strand RNA genome as a template to generate negatively stranded RNAs that will serve as templates for additional positive-strand mRNAs. During negative-strand transcription, RTCs will occasionally encounter a body transcription regulatory sequence (TRS-B). These occur at various positions within the viral genome. These sequences work in combination with a single leader transcription regulatory sequence (TRS-L) at the 5’ end of the genome. When RTCs encounter a TRS-B sequence, they can “jump” to the TRS-L sequence, thereby generating a negative-strand RNA that omits a large portion of the genome (Figure 3) (Sola et al., 2015). These negative-strand RNAs then serve as templates for transcription of positive-strand subgenomic mRNAs.. Positive-strand subgenomic RNAs created by this mechanism now have downstream AUGs in an optimal start codon context, allowing ribosomes to initiate translation at locations normally not available in the full length genomic RNA. These subgenomic RNAs serve as mRNAs for viral proteins encoded downstream in the genome.

**Figure 3:**
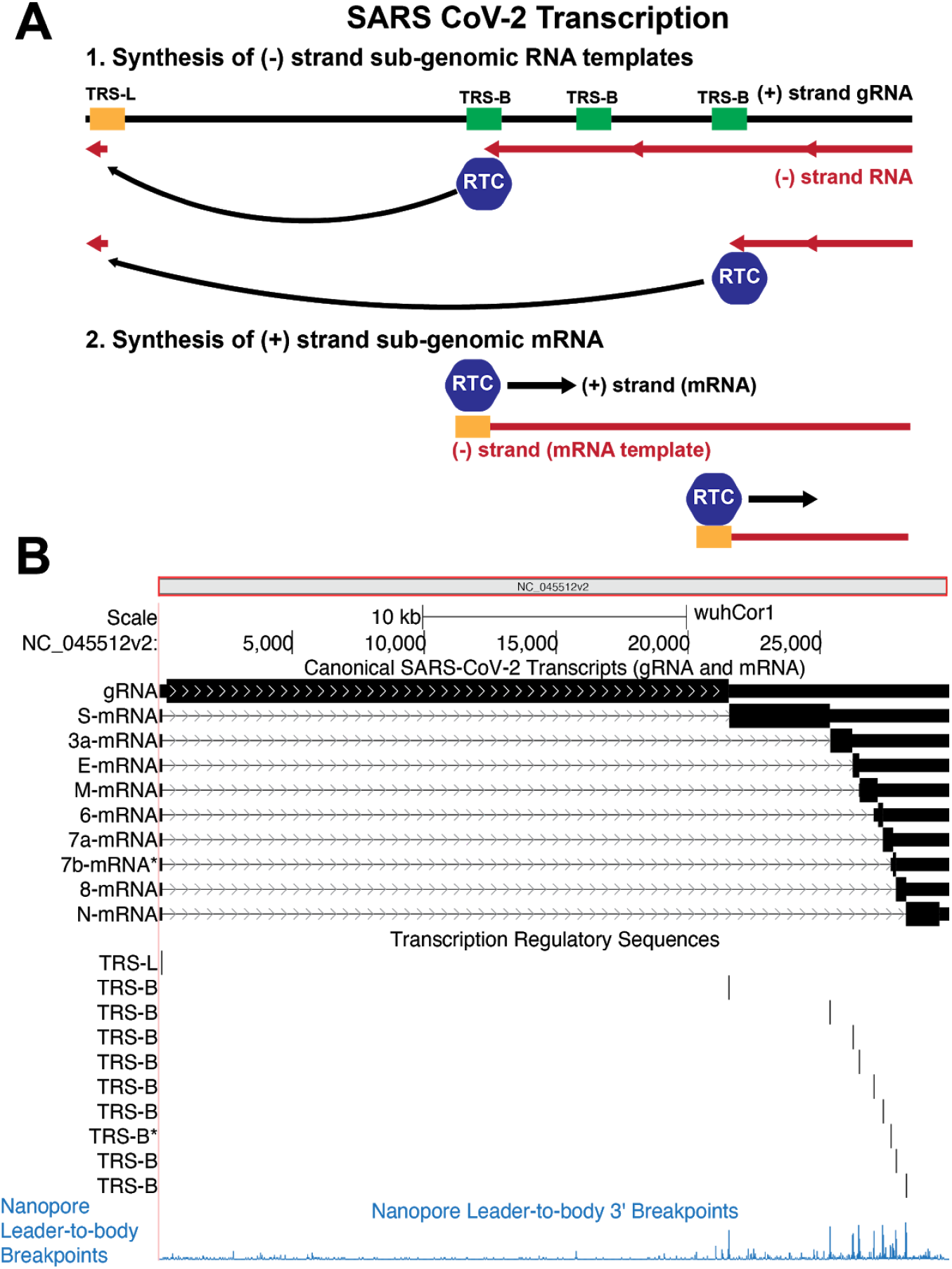
Molecular and Genomic Visualizations of SARS-CoV-2 Transcription on the browser. A) SARS-CoV-2 produces mRNAs by discontinuous transcription. First replication-transcription complexes (RTCs) initiate generation of (−) RNA strand (red strand) from the positive strand (black). When RTCs encounter a body transcription regulatory sequence (TRS-B) they have a chance to “jump” to the TRS-Leader (TRS-L) via long range RNA-RNA interactions. Alternatively they can proceed as usual transcribing along the genome. The jumping process generates several different species of (−) RNA strands that lack sequence between the various TRS-B sequences and the TRS-L. These (−) strands then serve as templates for positive transcription from the TRS-L to generate a variety of viral mRNAs that produce different viral proteins. B) A simple, compact and machine-readable genomic visualization for this complex biological process. (Top) All viral mRNA species are shown as annotations on the reference genome. Black bars represent nucleotides present in an mRNA species while arrows represent the sequence that has been skipped during discontinuous transcription. Thick black bars represent the coding sequence predicted to be translated in these RNA species. (Middle) The core TRS motif, ACGAAC, annotated on the genome, corresponds to transcript junctions. (Bottom) Experimental data representing breakpoints that are fusions of TRS-B to TRS-L sequence identified by Oxford Nanopore direct RNA sequencing (Kim et al., 2020). High peaks indicate that the 5’ TRS-L sequence is found directly upstream of the annotated bases in viral RNAs. The majority of these breakpoints correlate with TRS-B motifs. (Live Interactive Session: http://genome.ucsc.edu/s/SARS_CoV2/Figure3)

On the SARS-CoV-2 browser we have included a transcriptome track which annotates each predicted TRS site in the reference genome using the presence of the motif ACGAAC, the reported core TRS for SARS-CoV (Yount et al., 2006). The track also includes the canonical mRNA produced from the full length positive-strand RNA, as well as the sub-genomic mRNAs produced by TRS-B to TRS-L “jumps” that have been experimentally validated by transcriptomic sequencing (Kim et al., 2020). Although discontinuous transcription is mechanistically quite distinct from eukaryotic splicing, we annotate the “jump” from a TRS-B to the TRS-L in a manner analogous to introns, as this allows proper interpretation of the transcript. In addition, we have included annotations of several recently reported experimentally observed sub-genomic RNAs, some of which have non-canonical junctions (Kim et al., 2020). As the SARS-CoV-2 transcriptome continues to be elucidated, we will update and add appropriate tracks. (Note in press: we are currently adding similar data from (Nomburg et al., 2020)).

### The SARS-CoV-2 proteome

SARS-CoV-2 proteome consists of 2 polyproteins, 4 structural proteins, and possibly 9 accessory proteins (Brian & Baric, 2005; Fehr & Perlman, 2015; Wu et al., 2020). The two polyproteins are processed into 16 non-structural proteins, leading to as many as 29 proteins that should be considered when performing analyses (Figure 4).

**Figure 4:**
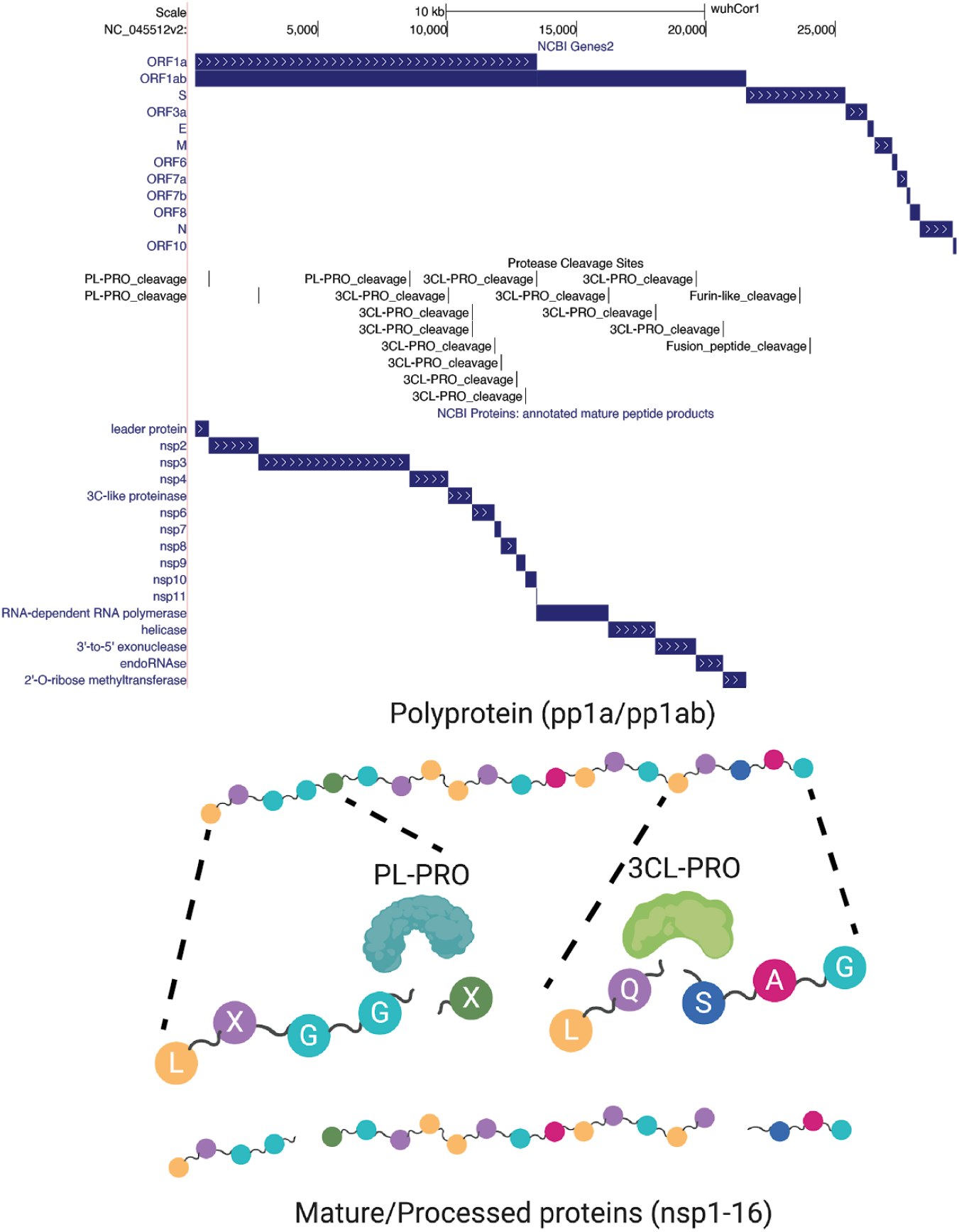
Cleavage of viral polyproteins. (Above) Browser track showing all annotated viral ORFs from the NCBI Genes track. Below the ORFs are the mature protein products that result from cleavage of the viral polyproteins by the viral proteases nsp3/PL-PRO and nsp5/3CL-PRO, as well as a track showing annotated cleavage sites. Also shown are two sites in the S (Spike) protein (furin_like_cleavage and fusion_peptide_cleavage) that are recognized by host cellular proteases instead of the above two viral proteases. Cleavage of coronavirus Spike protein generates mature subunits that allow the virus to enter cells. (Below) Cartoon representation showing abstractly the cleavage of polyprotein peptide sequences by the viral proteases to generate mature proteins. The viral polyproteins are cleaved by the PL-PRO protease at 3 locations that match the amino acid pattern LXGGX (X = any amino acid) as indicated; 3CL-PRO cleaves many more sites, typically at the pattern LQSAG as shown. (Live Interactive Session: http://genome.ucsc.edu/s/SARS_CoV2/Figure4)

Proteins pp1a and pp1ab are polyproteins that are products of ORF1a and ORF1ab respectively which are both produced by translation of the full length genomic RNA. To generate pp1a the ribosome initiates translation at the AUG at nucleotides 266-268. Translation continues in the canonical manner until a UAA stop codon is encountered at nucleotides 13,481-13,483. That produces the 4405 amino acid pp1a polyprotein. This polyprotein contains within itself two viral proteases (nsp3/PL-PRO) and (nsp5/3CL-PRO) that cleave the pp1a polyprotein into 11 mature non-structural proteins (nsp1-11, see Figure 4) (Barretto et al., 2005; Zhang et al., 2020)(Barretto et al., 2005).

The pp1ab polyprotein is generated by initiating translation at the same start codon as pp1a; however, the viral RNA contains within it a “slippery” sequence and structured RNA element that occasionally causes the translation machinery to slip near nucleotide C13468 and therefore read this nucleotide twice: once as the final nucleotide in the AAC codon for amino acid N4401, and once again as the first nucleotide in the CGG codon for amino acid R4402) (Bekaert & Rousset, 2005; Plant & Dinman, 2006). The result is a “programmed” −1 frameshift that means the pp1a “stop” codon is no longer in frame, lengthening pp1ab by 2695 entirely different amino acids. These additional amino acids encode essential non-structural proteins, (Figures 3 and 4), meaning that pp1ab encodes nsp1-10 as well as nsp12-16. In order to properly display this frameshift on the genome browser, we use specialized colored codons that lead to faithful translation of each ORF (Figure 5).

**Figure 5:**
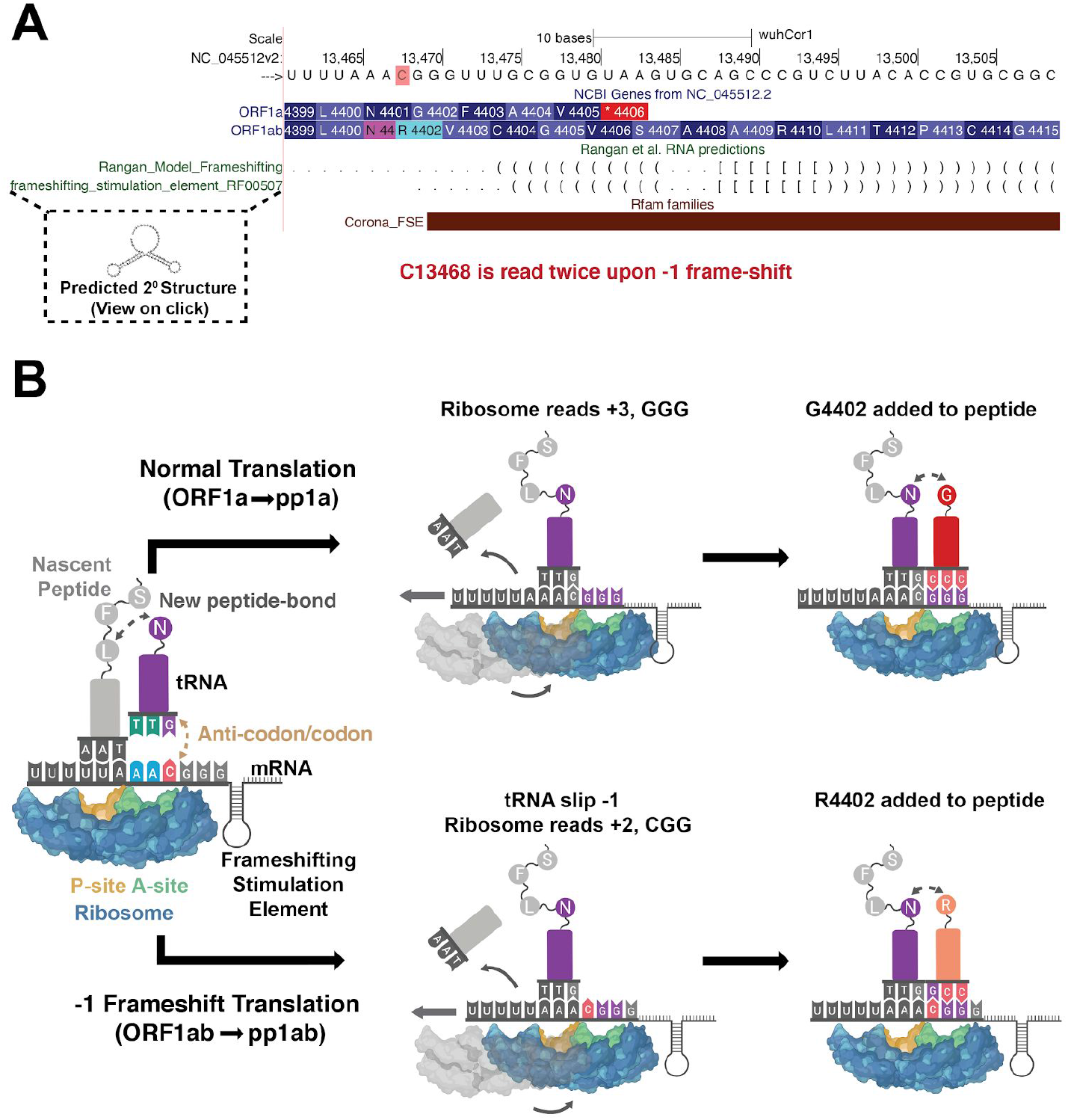
Orf1a/Orf1ab Ribosomal Frameshifting. A) Genome Browser annotations detailing translation of orf1a and orf1ab via ribosomal frameshifting. The red highlighted C is read twice by the ribosome due to the upstream poly-U tract and downstream frameshifting RNA structure annotated in the RFAM and RNA predictions track. Predicted base pairing reported by Ragan et al., 2020 and putative secondary structures are visible upon clicking on annotations. Note that tertiary interactions are not shown. (Live Interactive Session: http://genome.ucsc.edu/s/SARS_CoV2/Figure5) B) Schematic representation of ribosomal frameshifting to generate distinct protein products from ORF1a and ORF1ab. After the AAC in the A site of the ribosome recognizes its cognate tRNA, N4401 is added to the nascent peptide and the ribosome prepares to move N4401 to the P site of the ribosome and read the next codon at the A site of the ribosome. Normally this will occur canonically (above) with the ribosome moving +3 nucleotides on the mRNA (GGG) leading to addition of G4402 and normal translation of pp1a. However, occasionally (~10% of the time in SARS-CoV reporter constructs (Plant & Dinman, 2006)) the ribosome will slip due to the highly structured frameshifting element (depicted here as a simple cartoon stem loop) with the bound tRNAs slipping −1 nucleotide, and causing C13468 to remain in the A site of the ribosome. This results in +2 movement along the mRNA and an overall −1 frameshift. Therefore the next codon read is CGG and R4402 is added. Since the pp1a stop codon at 4406 is no longer in frame, ~2700 additional amino acids encoding nsp12-16 are added to the polyprotein.

The remaining proteins are produced through traditional ribosomal recognition of AUG sequences on sub-genomic RNAs as shown in Figure 3. Of particular interest is the Spike protein (S), which is the target of most immune-based therapies, governs the entry of the virus into the cell and is cleaved by host proteases as described in Figure 4. Other therapeutic targets include the RNA-dependent RNA polymerase (RdRp) protein that makes copies of the viral RNA genome (also known as Pol/nsp12), the virally encoded proteases 3CL-PRO and PL-PRO, the viral envelope protein E, and membrane protein M, and the nucleocapsid protein N, which organizes the RNA genome of the virus in viral particles.

SARS-CoV-2 has at least 5 accessory proteins encoded by ORF3a, ORF6, ORF7a, ORF7b, and ORF8. Additional accessory ORFs, not in the NCBI and Uniprot SARS-CoV-2 annotations, have been observed in other coronaviruses (ORF3b, ORF9b, ORF9c), although it is not clear whether these ORFs produce functional proteins in SARS-CoV-2 (Gordon et al., 2020; Schaecher & Pekosz, 2010). Additionally, ORF10, present in most annotation sets, has little experimental support as a protein-coding gene. (Davidson et al., 2020; Kim et al., 2020). A variety of variant ORFs have also been recently reported by non-canonical sub-genomic mRNAs (Nomburg et al., 2020).

### An overview of SARS-CoV-2 genome annotation tracks

We provide several standard annotation tracks based on molecular data generated by experimental and computational analyses of the SARS-CoV-2 genome. These annotations are sorted into groups as described below.

#### Mapping and Sequencing

Tracks in this group are all based on short segments of local nucleotide composition. For instance, users can immediately analyze inherent features of the sequence such as GC-content, and DNA restriction enzyme recognition sites. We have also added tracks that are specific for this viral genome: Covid-19 qPCR primers from WHO-listed detection kits (*Open COVID-19*, 2020), Nanopore sequencing primers from the ARTIC Network (*artic-ncov*, 2019) and high-scoring and validated Cas13 CRISPR guides (Abbott et al., 2020; Wessels et al., n.d.). The Crowd-sourced annotations track (see below) contains CRISPR guides used in SARS-CoV-2 detection via Cas12 (Broughton et al., 2020), and Cas13 (Metsky et al., 2020; Wessels et al., 2020), as well as LAMP primers from various sources, e.g. (Park et al., 2020). These tracks can be used in combination with the Variants tracks to determine if specific primers or detection methods might be less effective at detecting certain viral clades (Figure 6).

**Figure 6:**
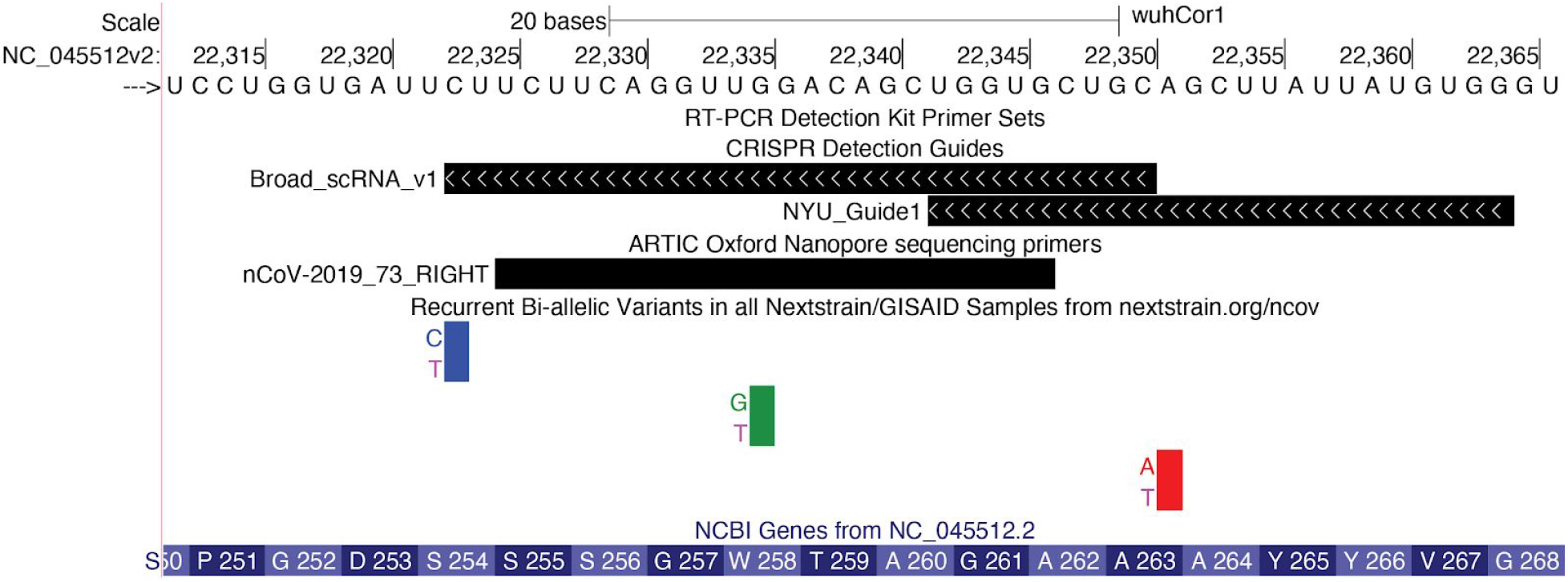
Variation in regions covered by CRISPR guides and PCR primers. Browser view of a portion of the viral genome coding for the S protein. CRISPR guide sequences for two Cas13-based detection kits developed by the Broad Institute (Metsky et al., 2020) and NYU are visible. Also visible is the right (3’ end) primer of primer-pair number 73 for whole genome assembly using the nanopore protocol from the ARTIC network Version 3. Although the Variants track reports three mutations in this region, non-reference alleles have been observed only 7 (C>T), 2(G>T), and 2(A>T) times in 4353 sequences (observed when mousing over, data not shown), suggesting that these regions are reasonable targets for primers and guides. The 7 instances of C>T are not alarming. An excess of C>T mutations from sequencing is observed throughout the viral genome, and is likely due to spontaneous deamination of cytosine into uracil or APOBEC RNA editing (Simmonds, 2020) (Live Interactive Session: http://genome.ucsc.edu/s/SARS_CoV2/Figure6)

#### Genes and Gene Predictions

These tracks contain information centered around the genes in the viral genome, as illustrated in figures above. For instance, the NCBI Gene track contains annotations of viral gene models from NCBI. As viral genes often have many names (e.g. nsp12/RdRp/Pol), many of these tracks list synonyms or notes in additional fields (viewable upon clicking an annotation) so that researchers can compare the annotations with the nomenclature that they are most familiar with. Additional tracks contain information such as interactions between viral proteins and human proteins from affinity-purification and mass-spectrometry experiments (Gordon et al., 2020), PDB structures (Figure 7), and RFAM and other predicted RNA structure annotations (Rangan et al., 2020) (Figure 5).

**Figure 7:**
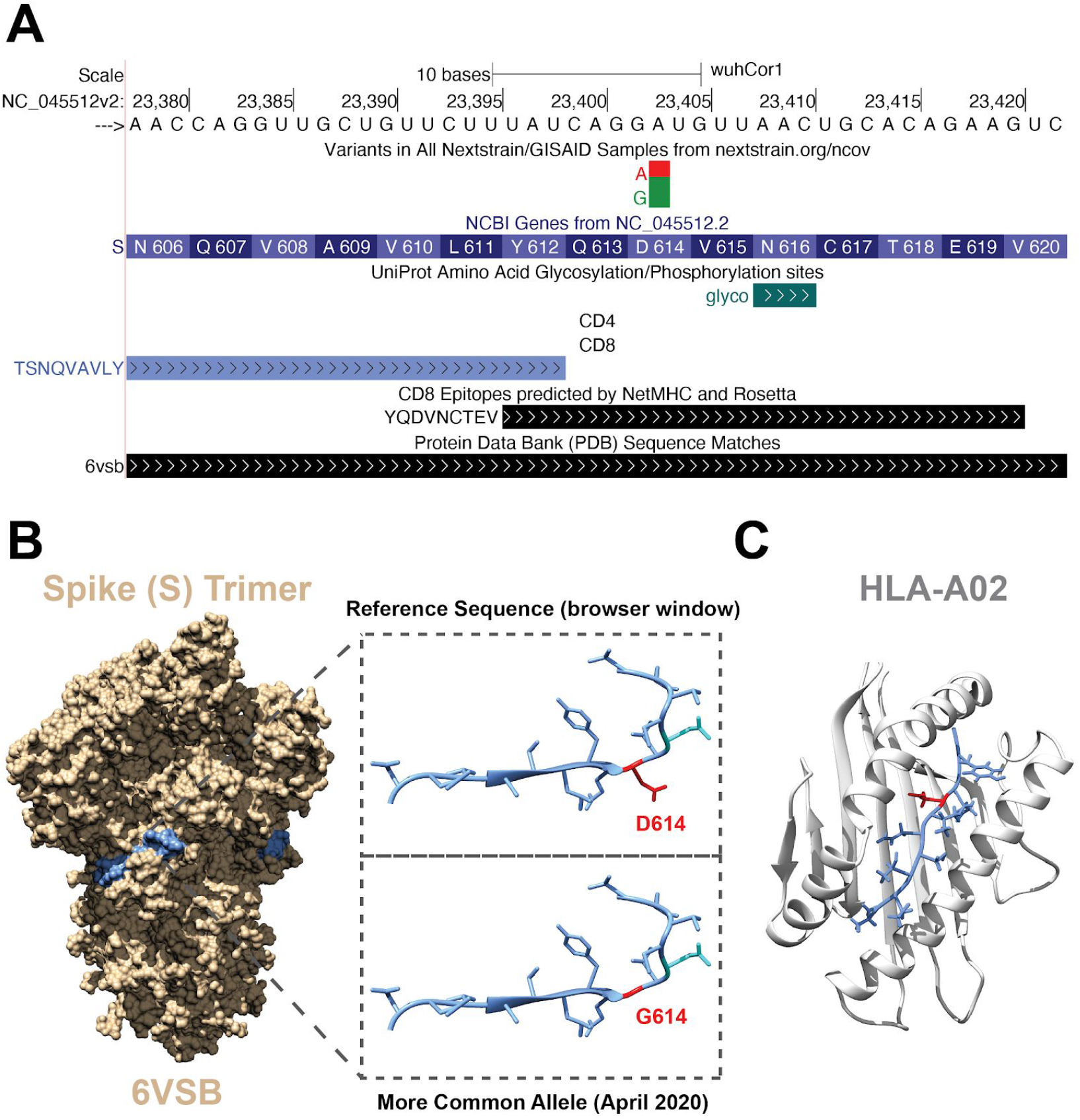
Combining Data Tracks to Generate Hypotheses. A) Browser view of a region of the viral genome that codes for part of the S (Spike) protein. The variants track shows an A>G mutation that causes the amino acid change D614G that is now found more commonly than the reference nucleotide from the original Wuhan outbreak (Andersen et al., 2020; Korber et al., 2020). Additional tracks display peptides within the virus that are predicted to be immunogenic. It is clear that the D614G mutation is contained within a predicted immunogenic peptide. Also shown is an annotated glycosylation site at amino acid 616 (highlighted in aqua) which can affect epitope recognition. B) Structure of the Spike (PDB ID: 6VSB) trimer. Highlighted in blue is the amino acid sequence viewed in A). Inset shows a close up view of the blue region with amino acid side chains. Highlighted in red is (Top) D614 (the product of the allele present in the original reference genome) and (Bottom) G614 (red) substituted in the structure using UCSF Chimera (Pettersen et al., 2004). C) Structure of the immunogenic peptide YQDVNCTEV in complex with HLA-A*02:01. D614 (red) is nestled within the binding groove, leading to a hypothesis that the G614 mutation may alter binding. Note that although browser-based comparisons of this data lend insight into possible models for the increased frequency of G614, further evolutionary and experimental analyses are required to make definitive statements about the functional consequences of this mutation. (Live Interactive Session: http://genome.ucsc.edu/s/SARS_CoV2/Figure7)

#### UniProt Protein Annotations

Protein annotations from SwissProt/UniProt (UniProt Consortium, 2019) are an essential complement to the NCBI RefSeq gene annotations (Pruitt et al., 2007). These tracks display a variety of protein annotation data including: special regions highlighted by SwissProt curators (e.g. the region of the S protein that binds the human receptor protein ACE2, see Figure 8), the mature proteins that result from polypeptide cleavage (Figure 4), as well as protein domains and sites of post-translational modification, such as glycosylation (Figure 6) and phosphorylation sites.

**Figure 8:**
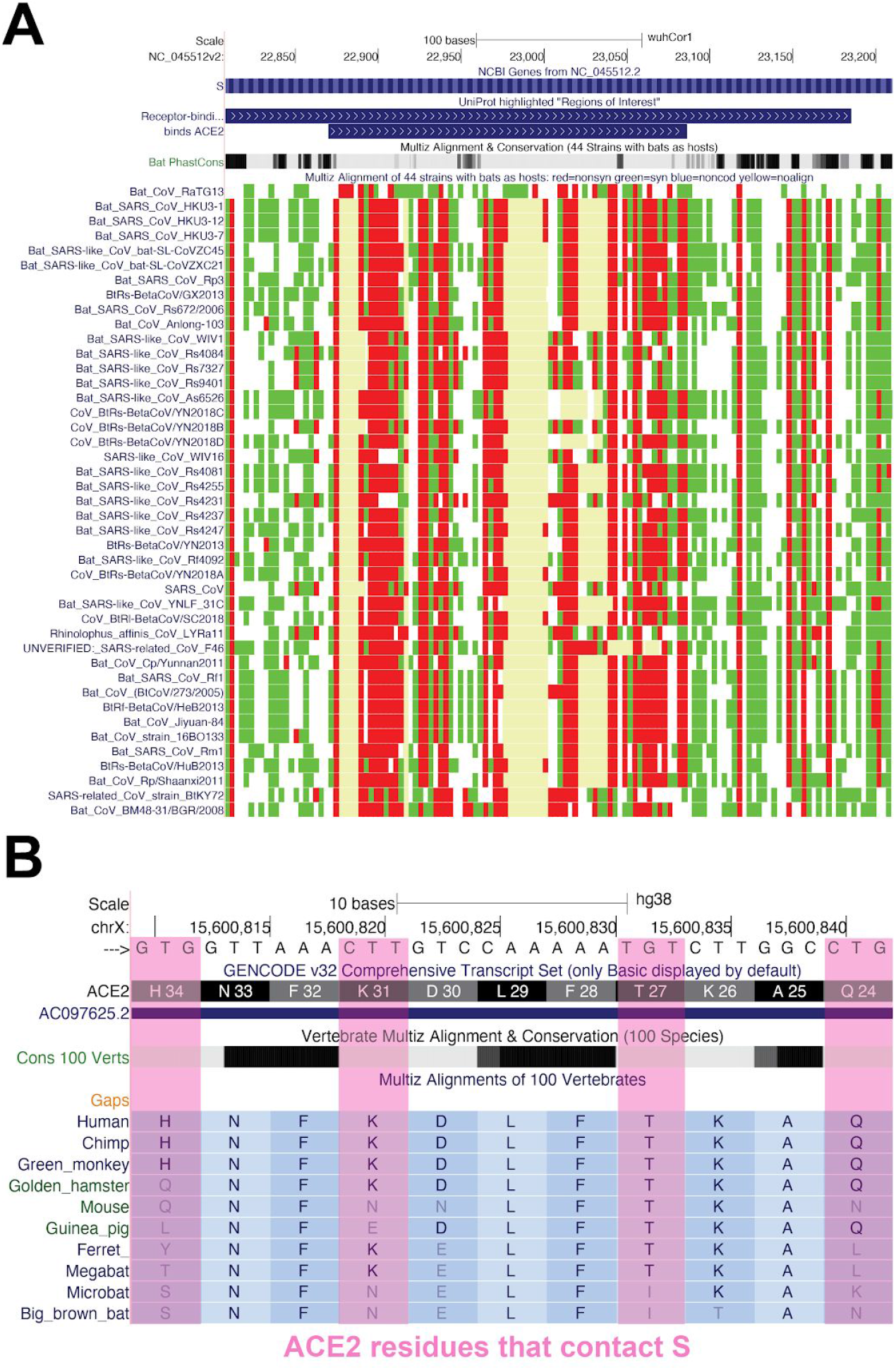
Comparative Genomic Analyses of SARS-like Coronaviruses. A view of the “44 Bat CoVs” track in the ACE2 binding region of the Spike protein. Red indicates a nonsynonymous change, green a synonymous change, and pale yellow (occasionally with blue border) indicates regions where no alignable sequence exists. The high divergence (red and pale yellow) in the amino acids within the ACE2 binding site (upper track) relative to those outside of the binding site is an indication of positive selection within the binding site. The “Bat PhastCons” track immediately above the multiple alignment summarizes per-base evolutionary rates for the nucleotide positions in the virus, light gray regions are more rapidly evolving (less conserved) than the black (very conserved) regions. This region of S is expected to experience selection as the ACE2 protein itself rapidly evolves between species in a “genetic arms race” with viruses that use this site. (Live Interactive Session: http://genome.ucsc.edu/s/SARS_CoV2/Figure8a) B) Human Genome Browser view of residues known to contact (in human) coronavirus spike proteins (pink) aligned to a variety of other species. These residues are more rapidly evolving (less conserved) in vertebrates (gray bars) than those that do not contact the Spike (solid black bars indicating strong vertebrate conservation). (Live Interactive Session: http://genome.ucsc.edu/s/SARS_CoV2/Figure8b)

#### Immunology

These tracks contain SARS-CoV-2 protein epitopes from publications. Included are epitopes that are predicted and/or validated to be immunogenic (Figure 7). The data features both linear epitopes recognized by B-cell receptors or antibodies as well as information on conformational epitopes (Pinto et al., 2020; Yuan et al., 2020) (recorded in Crowd-sourced annotations track, data not shown). These tracks also display epitopes recognized by CD8-positive or CD4-positive T-cells when presented by HLA molecules on host cells (Grifoni et al., 2020a; Poran et al., 2020). Where possible, the latter are organized according to the HLA allele of the host used in their presentation. Note that the four structural proteins, Spike (S) protein, the Membrane (M) protein, Envelope (E), and the Nucleocapsid protein (N) are the most frequent targets of the host immune system (Fast et al., 2020; Grifoni et al., 2020b) and many of these datasets are restricted to one of these regions, usually the Spike protein. For the track “CD8 RosettaMHC”, interactive 3D models from Rosetta are available after clicking on the annotated epitope (Nerli & Sgourakis, 2020). We will continue to update this track group as validation and identification of epitopes continues.

#### Comparative Genomics

This group contains two tracks that show multiple alignments built from sequences provided by NCBI/INSDC: 1) “119 Vertebrate CoVs”, an alignment of 119 sequences, most of which are human coronaviruses (though not necessarily SARS-CoV-2), with various animal viruses and 2) “44 Bat CoVs”, an alignment of 44 various bat coronaviruses most closely related to human SARS CoV-2. The mutations in the alignments are colored by their effect on the protein (white: no difference, red:nonsynonymous, green:synonymous, blue:noncoding, yellow:missing data due to unalignable, absent or unknown sequence). Analysis of evolutionary rates derived from comparative genomics, including insertions and deletions, can pinpoint functionally interesting sections of the viral genome. For instance, comparisons of SARS-CoV-2 with less pathogenic coronaviruses suggest features of the genome that may be related to increased pathogenicity (Gussow et al., 2020). Features identified in that analysis currently appear in the Crowd-sourced annotations track, but will eventually be displayed in the Comparative Genomics group. Similarly, comparison of coronaviruses across species and across host genomes can clearly illustrate evolutionary features such as accelerated evolution at receptor-spike binding regions (Figure 8) (Demogines et al., 2012).

#### Variant and Repeats

These tracks contain information on the variation and evolutionary patterns observed in SARS-CoV-2 sequences from different samples taken over the world. “NextstrainVars” (Figure 9) contains the time-stamped molecular phylogenetic tree produced by the Nextstrain team (Hadfield et al., 2018) based on complete and quality-controlled viral genomes from GISAID (Shu & McCauley, 2017). The tree is shown in pack and squish views. Dense and full mode display the frequencies at which these variants are found (data not shown). In the tree, samples are sorted according to their order in the JSON file produced by Nextstrain describing the phylogenetic tree, and appear in different colors according to clades identified by Nextstrain. Because data were collected by GISAID from labs all over the world the mutation calls do contain occasional errors. To avoid noise in the display, tools are provided to filter these data to show only the well-supported mutation calls. Additional filters allow users to set thresholds for minor allele frequency and to display data for specific clades.

**Figure 9:**
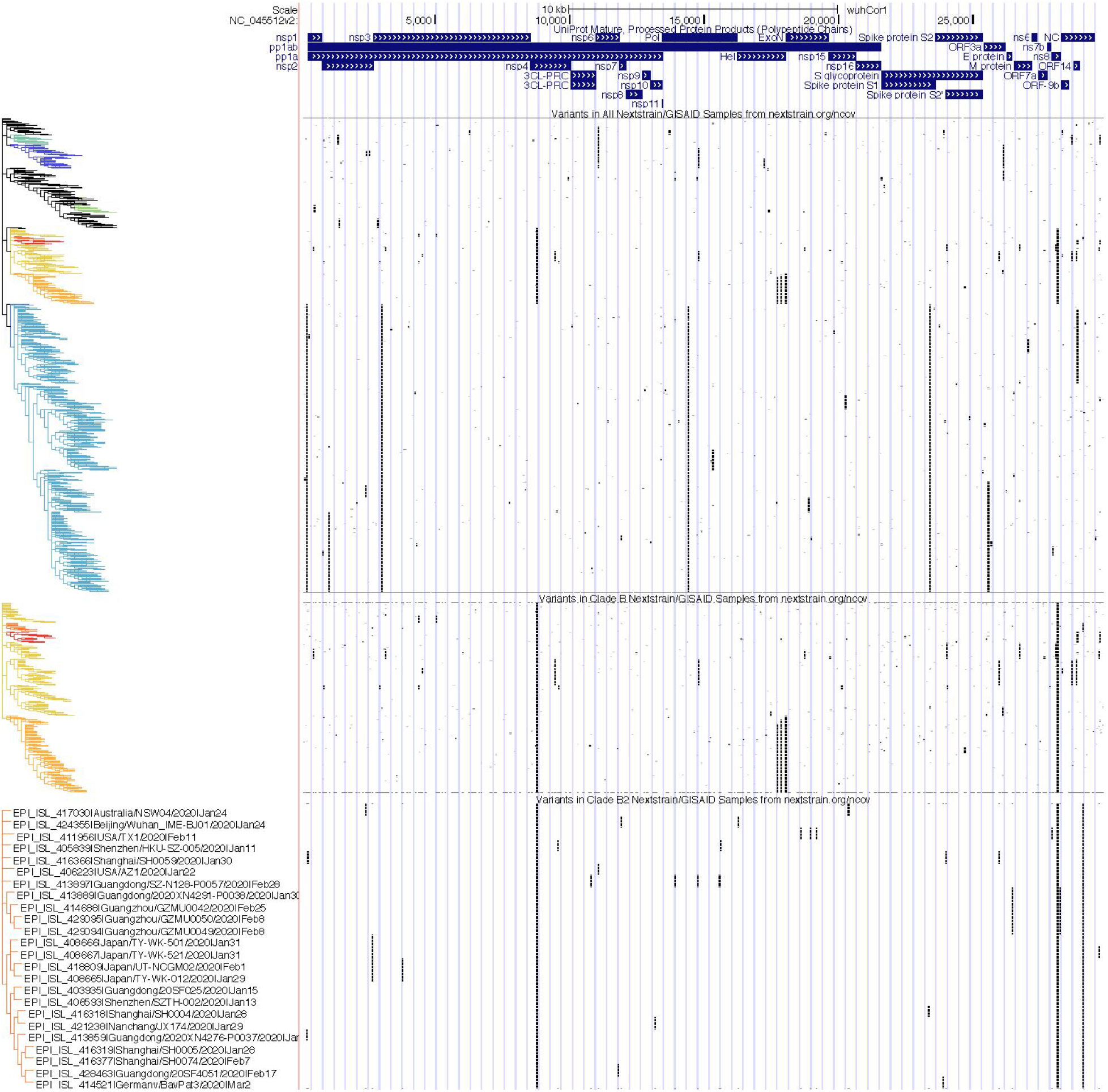
Phylogenetic Analyses of Clade-Specific Variation in SARS-CoV-2. SARS-CoV-2 mature protein products are shown at the top of the display for context. Below that are three tracks containing Nextstrain phylogenetic trees and clades for selected SARS-CoV-2 genomes sequenced from samples deposited in GISAID. Each row represents a single viral genome, black bars represent the presence of a mutation in the genome submitted to GISAID compared to the reference at that genomic position. As expected, many mutational patterns cluster with branches of the phylogenetic tree. The first track contains data from all 4147 sequences available from Nextstrain as of April 30, 2020. Clades identified by Nextstrain are colored by the same scheme used on the <u>Nextstrain site</u>. The middle track shows only samples from the Nextstrain B clade (warm colors) with increased vertical resolution, and the bottom track shows only samples from the Nextstrain B2 subclade of B (red-orange color) with even greater vertical resolution. At this resolution the sample identifier and additional sample information become visible, including time and location of the sample collection. (Live Interactive Session: http://genome.ucsc.edu/s/SARS_CoV2/Figure9)

#### Custom tracks

Users can add annotations for their own use or to share with other groups by clicking the “Custom tracks” button. Annotation data can be pasted directly into a text box (e.g. entering “NC_045512v2 509 515 binding-site” and clicking “Submit” will show an annotation at the nucleotide position 510-515 with the label “binding-site”) or annotations can be uploaded in a standard genomic file format. To share a custom track, users can create a link via the menu “My Data > My Sessions” and share this link directly. Session links can be shared multiple times and will always load the data exactly as originally saved.

#### Track Hubs

Track Hubs are similar to Custom Tracks but give the user more control over the display and make managing large collections of tracks easier (Raney et al., 2014). The files are hosted on a user’s web server and must be in binary formats such as BAM, VCF, bigBed or bigWig. The user defines the display parameters in configuration files (http://genome.ucsc.edu/goldenPath/help/hgTrackHubHelp.html). Track Hubs are displayed alongside the native annotation tracks for that assembly. They appear under the hub’s own track group on the main browser page. Track Hubs of sufficient general interest can be submitted for inclusion in our public hubs list at https://genome.ucsc.edu/cgi-bin/hgHubConnect#publicHubs.

#### Crowd-sourced data

To encourage shared community annotation of the viral genome without requiring that the submitter have detailed knowledge of genomic data type formats, a “Crowd-sourced annotations” track has been created. Its documentation includes a link (http://bit.ly/cov2annots) to a spreadsheet where anyone can enter the start and end position of some annotation, descriptive text and a link to a website or paper with the source information. A special link on the track description page adds a version of the track with all changes from today. Every night all new user annotations from the previous day undergo a brief manual check and are added to the public “Crowd-sourced annotations” track.

#### Data Access for Programmers

We maintain a public MySQL server (top menu bar “Downloads -> MySQL”) for accessing our annotation data and a JSON REST API (“Downloads -> JSON API”). We encourage data creators to contact us with requests for help with data conversion and about having their track hubs added to our list of public track hubs.

#### Outreach and Contact information

We offer free training for the Genome Browser over video calls. Outreach is supported by updates to the training documentation (https://genome.ucsc.edu/training/) with links to videos and in-depth descriptions of new Browser features. The training page includes information on how to submit a request for a webinar. General contact information for the UCSC Genome Browser can be found at https://genome.ucsc.edu/contacts.html, including information for accessing our email support list.

## Discussion

The rapid pace of SARS-CoV-2 research is generating a wealth of molecular and genomic data across a variety of databases. The UCSC Genome Browser is an established and highly-accessed web-based viewer and standardized repository of genomic data with extensive functionality and a 20 year track record of serving the genomics community. Here, we demonstrate the utility of this genomic viewing and downloading environment for the visualization and analysis of SARS-CoV-2 molecular data.

Currently, the SARS-CoV-2 browser has more than 50 tracks of various kinds of data, from gene annotations to phylogenetic trees of thousands of viral sequences. These data can be analyzed together in combinations chosen by the user to see named viral clades and keep track of viral strains as they emerge, gain easy access to both protein and RNA structures, survey predicted and validated immune epitopes, reveal mutational patterns in these immune epitopes, study glycosylation patterns that may affect antibody recognition, find locations of PCR primers and CRISPR guides and mutations that may affect them, compare SARS-CoV-2 to other viruses and analyze evolutionary patterns suggesting functional importance for particular regions, and to perform a variety of other analyses. We have carefully enhanced the display of certain annotations - such as the viral RNA’s ribosomal frameshift element - so that the browser correctly handles genomic features that are uncommon in most non-viral genomes, yet at the same time maintains reliable programmatic data access and automatic data updates.

The browser, and its underlying data organization, is a familiar environment for hundreds of thousands of biomedical researchers who use the human genome, allowing these scientists to download standardized data formats from a single source for custom analyses. Here we hope to also introduce these tools to virologists, epidemiologists, vaccinologists, antiviral therapy developers, and those seeking to repurpose existing biomedical resources and therapies to combat the virus and its pathological effects. In order to leverage the expertise of scientists who lack familiarity with genomic analyses but possess expertise in other areas, we have developed a “Crowd-Sourced Data” track in which users can simply enter the coordinates and name of a feature. This annotation is then displayed and shared with others on the browser, where it will be viewed and compared to the many existing data tracks. Together these features make the SARS-CoV-2 browser a simple but powerful tool for researchers to track developments in SARS-CoV-2 science, to detail the changes in the viral genome over the course of the pandemic, and to develop testable hypotheses and novel strategies to combat it.

As scientists continue to generate SARS-CoV-2 data, we will continue to rapidly process, display, and share these data on the SARS-CoV-2 browser. We urge authors to contact us at for help in properly citing, annotating and displaying their data in a clear, accurate, and intuitive manner on the browser so that it can reach the widest possible audience of researchers. Through this type of open collaboration, we believe the SARS-CoV-2 browser will facilitate the analysis and display of the collective molecular information needed to defeat the virus.

## Funding

The UCSC Human Genome Browser software, quality control, and training is funded by NHGRI, currently with grant 5U41HG002371-19. The SARS-CoV-2 genome browser and data annotation tracks are funded by generous individual donors including Pat & Rowland Rebele and a University of California Office of the President Emergency COVID-19 Research Seed Funding Grant R00RG2456.

## Conflict of interest statement

A.S.H., H.C, J.N.G., B.T.L., L.R.N., B.J.R., K.R.R., D.S., A.S.Z., W.J.K., D.H., and M.H. receive royalties from the sale of UCSC Genome Browser source code, LiftOver, GBiB, and GBiC licenses to commercial entities. W.J.K. owns Kent Informatics.

## Acknowledgements

We would like to thank Kord Kober and Kiley Charbonneau from the Kober lab at UCSF for their contributions to the crowd-sourced data track, as well as Holly Beale, Justin Sim, Alinne Gonzalez Armenta, Phil Berman, Nik Sgourakis and the rest of the scientists at UCSC and across the world for making these tracks of molecular information possible.

## Notes

https://genome.ucsc.edu/cgi-bin/hgTracks?db=wuhCor1

https://genome.ucsc.edu/covid19.html

